# Elevated activity of the mesolimbic dopamine system promotes feeding during pregnancy in mice

**DOI:** 10.1101/2025.10.21.683702

**Authors:** Tanya Pattnaik, Benjamin Wang, Patrick Sweeney

**Affiliations:** Department of Molecular and Integrative Physiology, University of Illinois Urbana-Champaign, Urbana, Illinois, USA; Neuroscience Program, University of Illinois Urbana-Champaign, Urbana, Illinois, USA

## Abstract

The pregnancy period is accompanied by increased feeding behavior to accommodate the elevated energy demands associated with fetal growth and development. However, the underlying neural circuitry and molecular mechanisms mediating increased feeding during pregnancy are largely unknown. Here, we utilize a combination of fiber photometry, chemogenetics, and mouse behavioral assays to characterize altered feeding behavior during pregnancy in mice. We uncover that pregnancy increases the activity of the mesolimbic dopamine system during both homeostatic and hedonic feeding behavior in mice. VTA dopamine neurons are ultimately required for promoting increased hedonic feeding during pregnancy as inhibition of these cells selectively reduces acute high fat diet intake in pregnant mice. Further, pregnant mice exhibit increased sensitivity to food deprivation, an effect which requires activity of the mesolimbic dopamine system. Together, these findings provide a circuit basis mediating altered hedonic feeding behavior and sensitivity to negative energy balance during pregnancy in mice.

**Highlights:**

- VTA dopamine neurons show enhanced responsivity to palatable food during pregnancy
- VTA dopamine neurons show enhanced responsivity to negative energy balance during pregnancy
- Nucleus accumbens dopamine is increased during homeostatic and hedonic feeding in pregnant mice
- VTA dopamine neuron activity regulates hedonic eating and fast-induced refeeding in pregnant mice

## Introduction

Mammals increase food intake during pregnancy to accommodate the elevated energy demands associated with fetal development^1,2^. Ultimately, increased feeding during pregnancy leads to a state of positive energy balance, resulting in elevated fat storage, which provides readily available energy stores for the metabolically demanding process of nursing^1,2^. Although increased feeding during pregnancy is critical for promoting healthy fetal development and growth, impaired energy homeostasis during pregnancy is associated with an increased risk of both the mother and her children developing obesity and type 2 diabetes later in life^1–5^. However, the core cellular and molecular mechanisms leading to increased feeding during pregnancy are incompletely understood.

Feeding behavior is initiated by neural circuits located in the hypothalamus, midbrain (i.e. ventral tegmental area), and hindbrain^6,7^. Neurons in these regions integrate hormonal and neuroendocrine cues which signal changes in long-term energy stores (i.e. via leptin from fat and ghrelin from the stomach) to initiate food intake during conditions of energy deprivation^6,7^. Following food consumption, meal derived satiety signals and GI distention regulate hypothalamic, midbrain, and hindbrain circuits to terminate ongoing feeding behavior^8^. Together, these processes control energy homeostasis by matching energy intake to changes in energy expenditure (i.e. homeostatic feeding behavior). In addition to homeostatic feeding behavior, consumption of palatable food also occurs in the absence of hunger (i.e. hedonic feeding)^9,10^. Although homeostatic and hedonic feeding occur independently of each other, energy state influences the propensity to seek and consume palatable energy dense foods^10^. Thus, palatable food seeking and consumption is increased in energy deprived animals, while satiety signals reduce the propensity to consume palatable, energy dense foods^9–14^. While the underlying neural circuitry controlling homeostatic and hedonic feeding is extensively studied, the effect of pregnancy on these neural pathways is not well understood.

Among the neuronal pathways controlling feeding, dopaminergic pathways are essential for promoting food seeking and consumption^9,15^. Dopamine deficient mice are aphagic and die of starvation unless provided supplemental nutrition via a feeding tube, suggesting that dopamine promotes food seeking and/or consumption^15^. Consistent with this notion, dopamine neurons in the ventral tegmental area (VTA) are robustly activated during food seeking and consumption of palatable food and exert an important role in signaling the post-ingestive reward associated with the consumption of calories^9,13,16–18^. VTA dopamine neurons project throughout the brain, including to the nucleus accumbens, amygdala, and prefrontal cortex^9^. In particular, VTA dopamine projections to the nucleus accumbens (NAc; mesolimbic dopamine pathway) represent a critical neural circuit controlling reward seeking behavior, including the seeking and consumption of palatable food^9,10^. Although most data suggest that dopamine promotes food seeking and palatable food intake, pharmacological studies provide contrasting results. For example, many dopamine mimetics are potently anorexic (i.e. cocaine and amphetamines), suggesting an appetite suppressive effect of dopamine transmission^9,19,20^. Thus, the specific role of dopamine in feeding is debated, and the contribution of dopamine to feeding behavior during pregnancy is largely unknown. Here, we utilized a combination of mouse feeding assays, *in vivo* imaging of dopaminergic transmission, and chemogenetics to characterize the role of mesolimbic dopamine signaling in feeding behavior during pregnancy.

## Results

### Pregnant mice exhibit hyperphagia for both regular chow and high fat diet

Although pregnancy is known to increase food intake in rodents^1,2^, the specific effect of pregnancy on homeostatic feeding, hedonic feeding, and food seeking behavior is incompletely understood. To test if pregnant mice exhibit elevated hedonic food consumption in the minutes following access to palatable foods, we provided palatable peanut butter (PB) chips to non-pregnant and pregnant mice. Mice were provided with PB chips daily for ten minutes for three days prior to testing to habituate the mice to PB chips and prevent food neophobia. Pregnant mice consumed more PB chip in the minutes following PB chip presentation compared to non-pregnant animals, indicating that pregnant mice exhibit enhanced palatable food intake during acute presentation of palatable foods (**Fig. 1A**). Next, we quantified daily food intake in non-pregnant and pregnant mice during the consumption of regular chow diet or a 60% high fat diet. Consistent with prior reports^1,21^, pregnant mice consume approximately 25% more calories than age-matched non-pregnant mice when provided regular chow diet (**Fig. 1B**). To characterize hedonic feeding behavior in pregnant mice during longer periods of access to palatable high fat foods, we provided *ad libitum* access of a palatable high fat diet (60% fat) to non-pregnant and pregnant mice. As expected, both pregnant and non-pregnant mice significantly increased their caloric intake following 24-hour access to high fat diet (compared to regular chow intake; **Fig. 1B and 1C**). However, pregnant mice continued to consume significantly more calories than non-pregnant mice for three consecutive days following HFD access. When calorie intake was adjusted to account for the percent increase in calorie intake on HFD (**Fig. 1C**, vs baseline regular chow intake), pregnant and non-pregnant mice exhibited a similar level of hyperphagia when provided with high fat diet. Further, in both non-pregnant and pregnant mice, calorie intake is similarly reduced on the second and third day of HFD access, compared to the initial day of HFD access, indicating that pregnant mice appropriately adapt their voluntary food intake in response to long-term access to hypercaloric diets (**Fig. 1C**).

**Figure 1:**
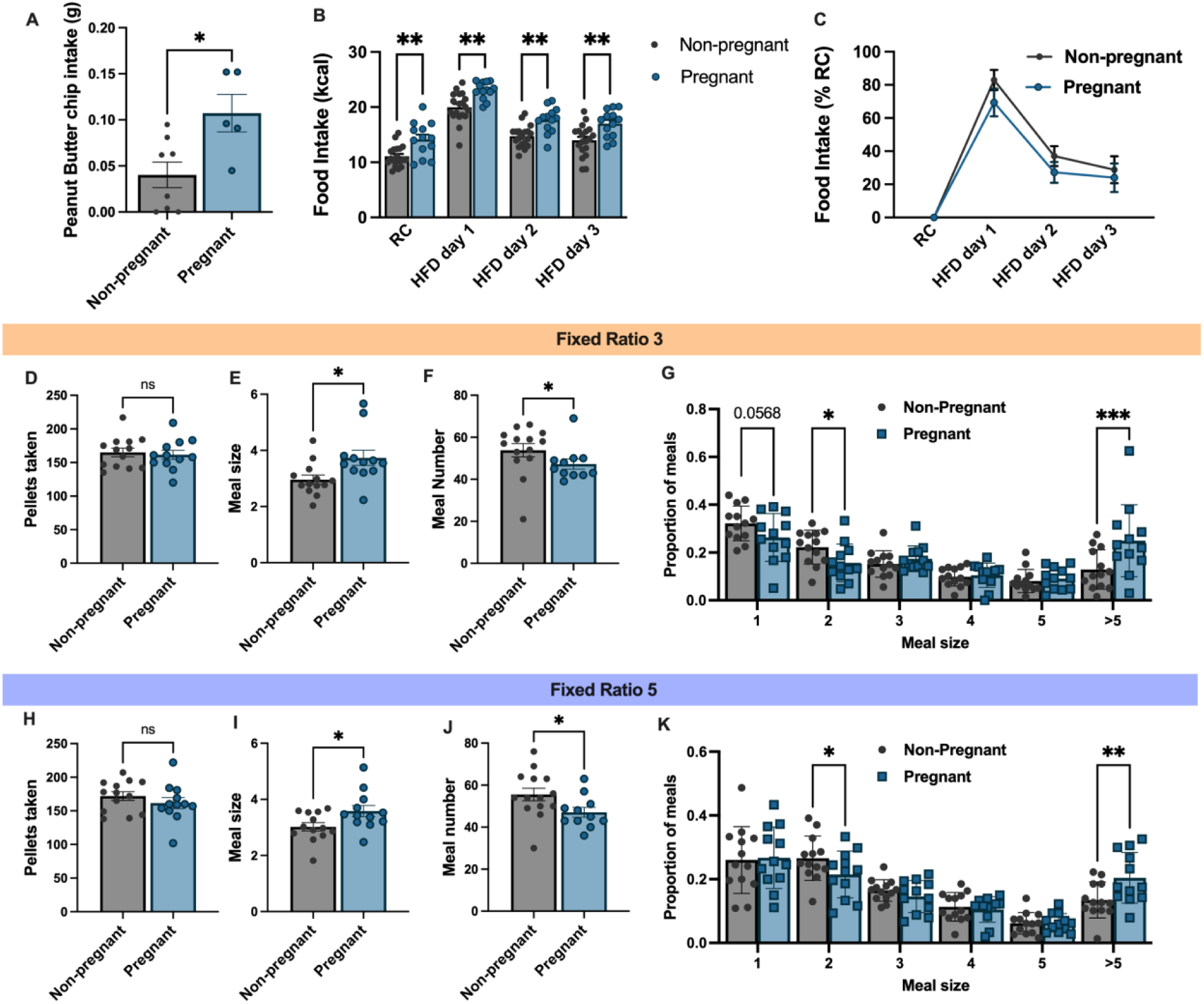
Pregnant mice exhibit hyperphagia for both regular chow and high fat diet. (A) Peanut butter chip intake in non-pregnant and pregnant mice during an acute 10-minute feeding assay. (B) Daily food intake (in kcal) of non-pregnant and pregnant mice (third trimester) fed a regular chow diet (RC) or a high fat diet (HFD). (C) Change in food intake upon access to high fat diet compared to regular chow food intake in non-pregnant and pregnant mice. (D-F) Pellets consumed (D), meal size (E), and number of meals (F) during fixed-ratio 3 (FR3) food seeking tasks in non-pregnant and pregnant mice. (G) Relationship between the percentage of meals consumed and the size of meals in non-pregnant and pregnant mice during FR3 feeding assays. (H-J) Pellets consumed (H), meal size (I), and number of meals (J) during fixed-ratio 5 (FR5) food seeking tasks in non-pregnant and pregnant mice. (K) Relationship between the percentage of meals consumed and the size of melas in non-pregnant and pregnant mice during FR5 food seeking tasks in non-pregnant and pregnant mice. Data points represent individual mice. Panels B, C, G, and k analyzed by 2-way ANOVA. Panels A, D, E, F, H, I, and J analyzed with unpaired Students t-test. ns (not significant), *p<0.05, **p<0.01, ***p<0.005.

### Pregnant mice consume larger meals in operant food seeking assays

Having characterized the feeding response to *ad libitum* access of regular chow or high fat diets, we next utilized home-cage operant feeding devices (feeding experimental device 3; FED3) to describe operant food seeking behavior in non-pregnant and pregnant mice^22^. Non-pregnant and pregnant mice were trained to nose poke on the left nose poke port to receive a single 20mg food pellet. Following successful training (greater than 70% of nose pokes occurring on the correct nose poke port), we quantified the meal size, meal frequency, and total pellets consumed over 24 hours in non-pregnant and pregnant mice during a fixed ratio 3 schedule of reinforcement (3 correct nose pokes leads to 1 20mg food pellet). In contrast to free access feeding conditions (**Fig. 1B**), pregnant mice did not consume more pellets than non-pregnant mice during FR3 feeding assays (**Fig. 1D**). There was, however, a significant shift in meal size and frequency in pregnant animals with pregnant mice consuming larger meals (**Fig. 1E**) but eating less frequently than non-pregnant mice (**Fig. 1F**). Pregnant mice exhibited a dramatic shift in the proportion of meals larger than 5 pellets, indicating that pregnant mice shift their meal structure towards larger, and more infrequent meals compared to non-pregnant animals (**Fig. 1G**). Similar findings were also observed during a fixed ratio 5 schedule of reinforcement (**Fig. 1H-K**). Therefore, pregnant mice exhibit altered feeding structure during operant food seeking assays, favoring larger and more infrequent meals compared to non-pregnant animals.

### VTA dopamine neurons exhibit enhanced responsivity to hedonic and homeostatic feeding in pregnant mice

Although hedonic feeding and meal structure is altered in pregnant mice (**Fig. 1**), the underlying neural circuitry mediating these changes is unknown. Given that ventral tegmental area (VTA) dopamine neurons modulate both hedonic feeding and meal size in mice^9,10,16^, we next utilized *in vivo* calcium imaging to characterize changes in VTA dopamine activity in pregnant mice. DAT-cre transgenic mice were injected with the genetically encoded calcium indicator GCAMP6s into the VTA and a fiber optic cannula was positioned into the VTA (**Fig. 2A**) to record changes in calcium activity as a proxy of neuronal activity. Following recovery from surgeries, mice were randomly separated into a pregnant group, which was mated with a male mouse, and a non-pregnant, age-matched control group (**Fig. 2B**). Given the critical role of VTA DA neurons in palatable food intake we first measured calcium activity in VTA DA neurons in response to the presentation of high fat diet (60% fat) in non-pregnant and pregnant mice. As expected, VTA DA activity increased in both non-pregnant and pregnant mice following consumption of HFD (**Fig. 2C**). The average increase in VTA DA activity was significantly larger in pregnant mice than in age matched non-pregnant animals during HFD consumption (**Fig. 2D**). However, pregnant and non-pregnant mice exhibited a similar maximum response in VTA DA activity following HFD presentation (**Fig. 2E**). The increased calcium signal in pregnant mice was not due to increased HFD intake in pregnant mice, as pregnant and non-pregnant mice consumed a similar amount of HFD during the brief ten-minute imaging session (**Extended Fig. 1**).

**Figure 2:**
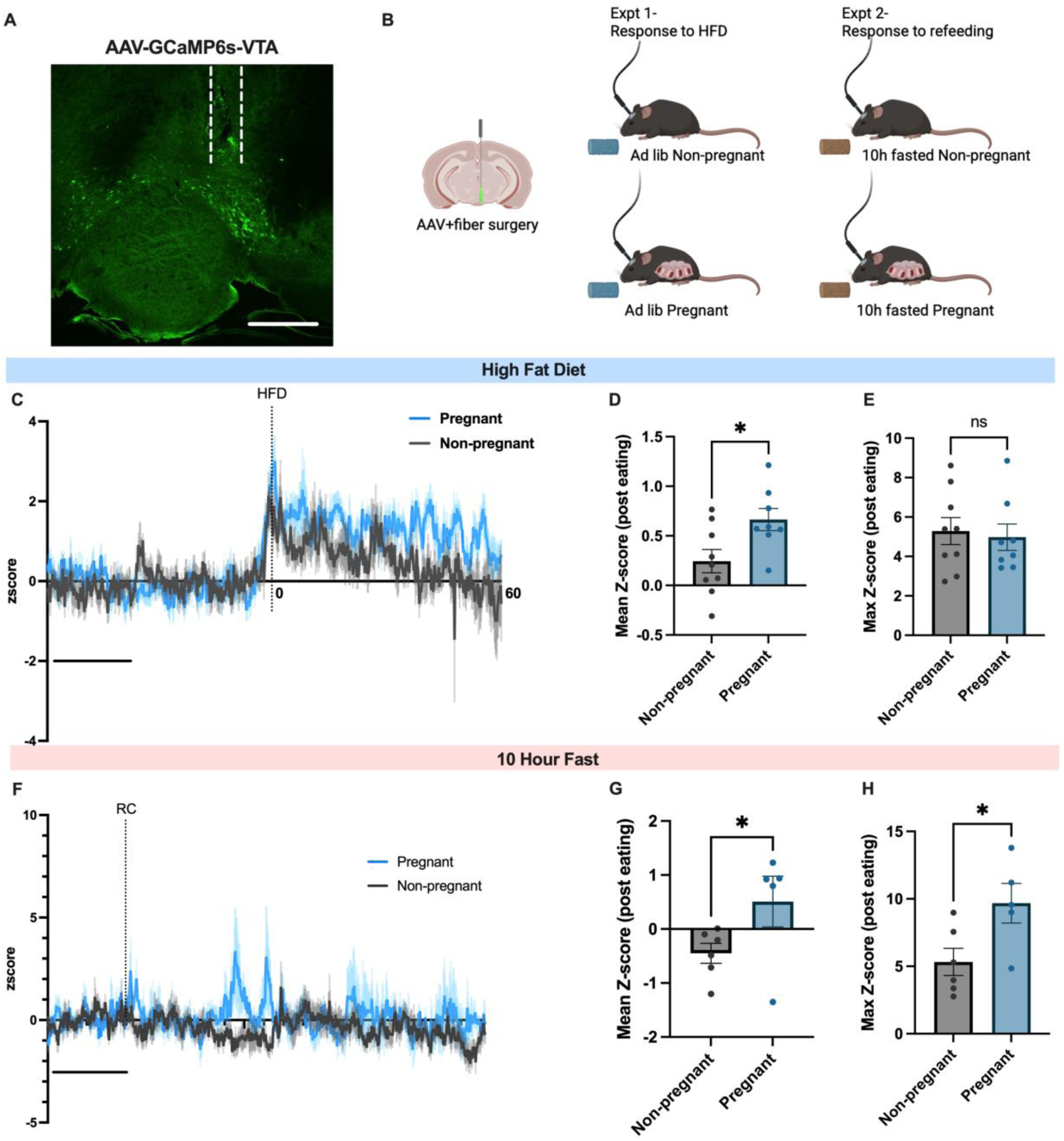
VTA dopamine neurons exhibit increased responsivity to homeostatic and hedonic feeding during pregnancy. (A) Representative image showing expression of GCaMP6s virus in VTA with fiber location. (B) Schematic showing experimental setup. (C) Average trace of the calcium signal in VTA dopamine neurons following the consumption of high fat diet. (D and E) Mean change in calcium signal (D) and maximum change (E) in VTA dopamine neurons following HFD consumption in non-pregnant and pregnant mice. (F) Average trace of the calcium signal in VTA dopamine neurons following the presentation of food after a ten hour fast in non-pregnant and pregnant mice. (G and H) Average change in calcium signal (G) and maximum change in calcium signal (H) following consumption of standard chow in non-pregnant and pregnant mice. Data in C and F represents average signal with standard error of the mean from all mice. Data points in D, E, G, and H represent individual mice. Data analyzed with unpaired Student’s t-test. ns (not significant), *p<0.05. Scale bar in A (500um), Scale bar in C (20 seconds), Scale bar in F (20 seconds).

In addition to promoting the consumption of palatable rewarding foods, VTA dopamine activity is also involved in promoting food seeking behaviors and reinforcement associated with food consumption in energy deprived mice^9,13,14,18^. Therefore, we next tested if the VTA DA response to standard food consumption differed between non-pregnant and pregnant mice. Both non-pregnant and pregnant mice were fasted for 10 hours and presented with familiar standard chow in their home cage while imaging the activity of VTA DA neurons with fiber photometry. The average signal intensity in VTA dopamine neurons in pregnant mice following refeeding was significantly higher than in non-pregnant mice (**Fig. 2F and G**). Further, the maximum increase in VTA DA activity was significantly higher in pregnant mice than non-pregnant animals following consumption of standard chow (**Fig. 2H**). These differences were not secondary to increased food consumption in pregnant mice as both non-pregnant and pregnant mice consume similar amounts of food during the acute testing session (**Extended Data Fig. 1**). Thus, VTA dopamine neuron activity is enhanced in pregnant mice during both hedonic and homeostatic feeding behavior (**Fig. 2**).

### Nucleus accumbens dopamine response is increased in pregnant mice during homeostatic and hedonic feeding

Although VTA dopamine neurons project to multiple downstream brain regions, the neuronal projection to the nucleus accumbens is particularly important for promoting hedonic feeding and food seeking behaviors^9,10,23^. Therefore, we next utilized genetically encoded dopamine sensors^24^ and fiber photometry to quantify the levels of dopamine transmission in the nucleus accumbens during homeostatic and hedonic feeding behaviors in non-pregnant and pregnant mice (**Fig. 3**). Genetically encoded dopamine sensors (GRAB-DA) were targeted to the nucleus accumbens (NAc), and a fiber optic cannula was positioned directly above the NAc to quantify changes in dopamine levels with fiber photometry (**Fig. 3A and B; Extended Data Fig. 2**). First, we measured changes in NAc dopamine levels prior to pregnancy and during the pregnancy period (or the equivalent time-period in non-pregnant control mice) after providing high fat diet to mice. The average increase in NAc dopamine levels following HFD consumption was significantly increased in pregnant mice, compared to baseline measurements prior to pregnancy (**Fig. 3C-D**). In contrast, the maximum increase in NAc dopamine levels was similar pre-pregnancy and during pregnancy in the minute following HFD consumption (**Fig. 3E**). Importantly, no difference in the NAc dopamine response to HFD consumption was observed in both time periods in non-pregnant control mice, indicating that these differences likely don’t result from order effects associated with repeated experiments (**Fig. 3I**). Further, a similar trend towards increased NAc dopamine levels was also observed in pregnant mice compared to non-pregnant animals tested on the same day (**Extended Data Fig. 3A**)

**Figure 3:**
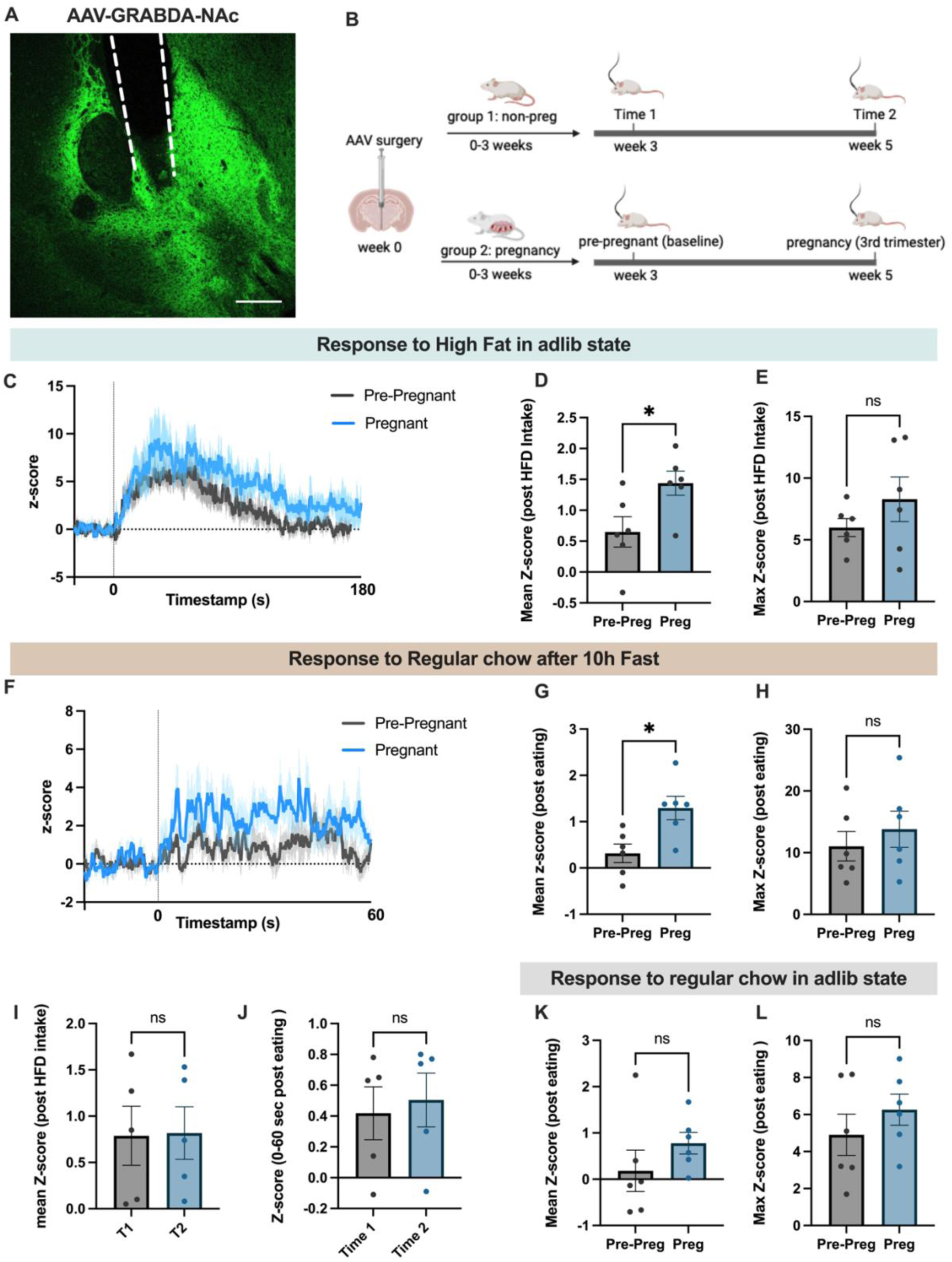
Dopamine levels in the nucleus accumbens exhibit increased responsivity to homeostatic and hedonic feeding during pregnancy. (A) Representative image showing expression of GRAB dopamine sensor in nucleus accumbens with fiber location in nucleus accumbens. (B) Timeline for experiments shown in figure 3. (C) Average trace of the nucleus accumbens dopamine signal following high fat diet consumption prior to pregnancy and during the pregnancy period. (D and E) Quantification of the data shown in C, comparing the mean Z-score following HFD consumption (D) and the maximum signal (E) following HFD consumption in non-pregnant and pregnant mice. (F) Average trace of the nucleus accumbens dopamine signal following food consumption after a ten hour fast before pregnancy and during the pregnancy period. (G and H) Quantification of the data shown in F, comparing the mean Z-score following food consumption (G) and the maximum Z-score following food consumption (H). (I and J) Average change in dopamine signal in non-pregnant mice at the two experimental timepoints shown in B following HFD consumption (I) or food intake following a ten hour fast (J). (K and L) Mean change in dopamine signal (K) and maximum change in dopamine signal (L) following the presentation of food to non-pregnant and pregnant mice. Mice were not food deprived prior to food presentation in panels K and L. Data points represent individual mice. All panels analyzed by paired Student’s t-test. ns (not significant), *p<0.05. Scale bar in A (300um).

Since we observed an increased VTA dopamine response to regular chow consumption in energy deprived mice during pregnancy (**Fig. 2**), we next measured changes in NAc dopamine levels pre-pregnancy and during the pregnancy period following food consumption after a ten hour fast (**Fig. 3F-H**). The average increase in nucleus accumbens dopamine levels was significantly greater in the 60 seconds following food consumption in pregnant mice compared to the pre-pregnancy period (**Fig. 3G**). However, mice exhibited a similar maximum change in NAc dopamine levels in the 60 seconds following food consumption during the pre-pregnancy period and the pregnancy period (**Fig. 3H**). No change in the average dopamine response was observed in the 60 seconds following food consumption in non-pregnant mice during the two recording sessions, indicating that these changes do not result from order effects associated with repeated imaging sessions (**Fig. 3J**). Furthermore, nucleus accumbens dopamine levels were also elevated in pregnant mice compared to non-pregnant control animals on the same testing day (**Extended Data Fig. 3B**), further validating increased NAc dopamine levels during food consumption in energy deprived pregnant mice.

To test if the increased NAc dopamine response during regular chow food consumption in pregnancy is specific to energy deprived mice we also monitored NAc dopamine levels in non-pregnant and pregnant mice following food presentation in *ad libitum* fed mice. In contrast to fasted animals, no difference in the NAc dopamine response was detected between non-pregnant and pregnant mice after food presentation in sated animals (**Fig. 3K-L**). Thus, pregnant animals specifically exhibit an increased mesolimbic dopamine response to negative energy balance compared to non-pregnant animals.

### VTA dopamine neurons regulate meal size and number in both pregnant and non-pregnant mice

Our prior *in vivo* imaging data (**Fig. 2 and Fig. 3**) indicate that pregnancy alters the activity of VTA DA neurons and dopaminergic transmission in the nucleus accumbens during homeostatic and hedonic feeding. We thus hypothesized that VTA dopamine neuron activity regulates feeding behavior during pregnancy in mice. To test this hypothesis, we targeted the chemogenetic inhibitor hM4Di or control virus expressing a fluorescent protein to VTA dopamine neurons in DAT-cre mice, and inhibited VTA dopamine neurons during feeding tasks in non-pregnant and pregnant mice (**Fig. 4A**). Feeding assays were performed using feeding experimental devices (FED3) to quantify changes in food intake, meal size, and meal number following inhibition of VTA dopamine neurons. CNO-mediated inhibition of VTA dopamine neurons did not alter the number of pellets consumed in non-pregnant or pregnant mice which were provided *ad libitum* access to food (**Fig. 4B**). As previously described (**Fig. 1**), pregnant mice consumed larger meals than non-pregnant mice (**Fig. 4C**). A similar increase in meal size was observed in pregnant mice compared to non-pregnant mice following either saline or CNO injections, indicating that VTA dopamine activity is not required for promoting increased meal size in pregnant mice (**Fig. 4C**). Inhibition of VTA dopamine neurons led to a similar increase in meal size in both non-pregnant and pregnant mice, and an equivalent decrease in meal number in both non-pregnant and pregnant mice (**Fig. 4C and D**). No differences between saline or CNO injections were observed for pellets consumed, meal size, or meal number in both non-pregnant and pregnant mice expressing control mCherry virus in VTA dopamine neurons, indicating no off-target effects of CNO or viral expression (**Extended Data Fig. 4**). Therefore, inhibition of VTA dopamine neurons increases meal size and reduces meal number to a similar extent in both non-pregnant and pregnant mice.

**Figure 4:**
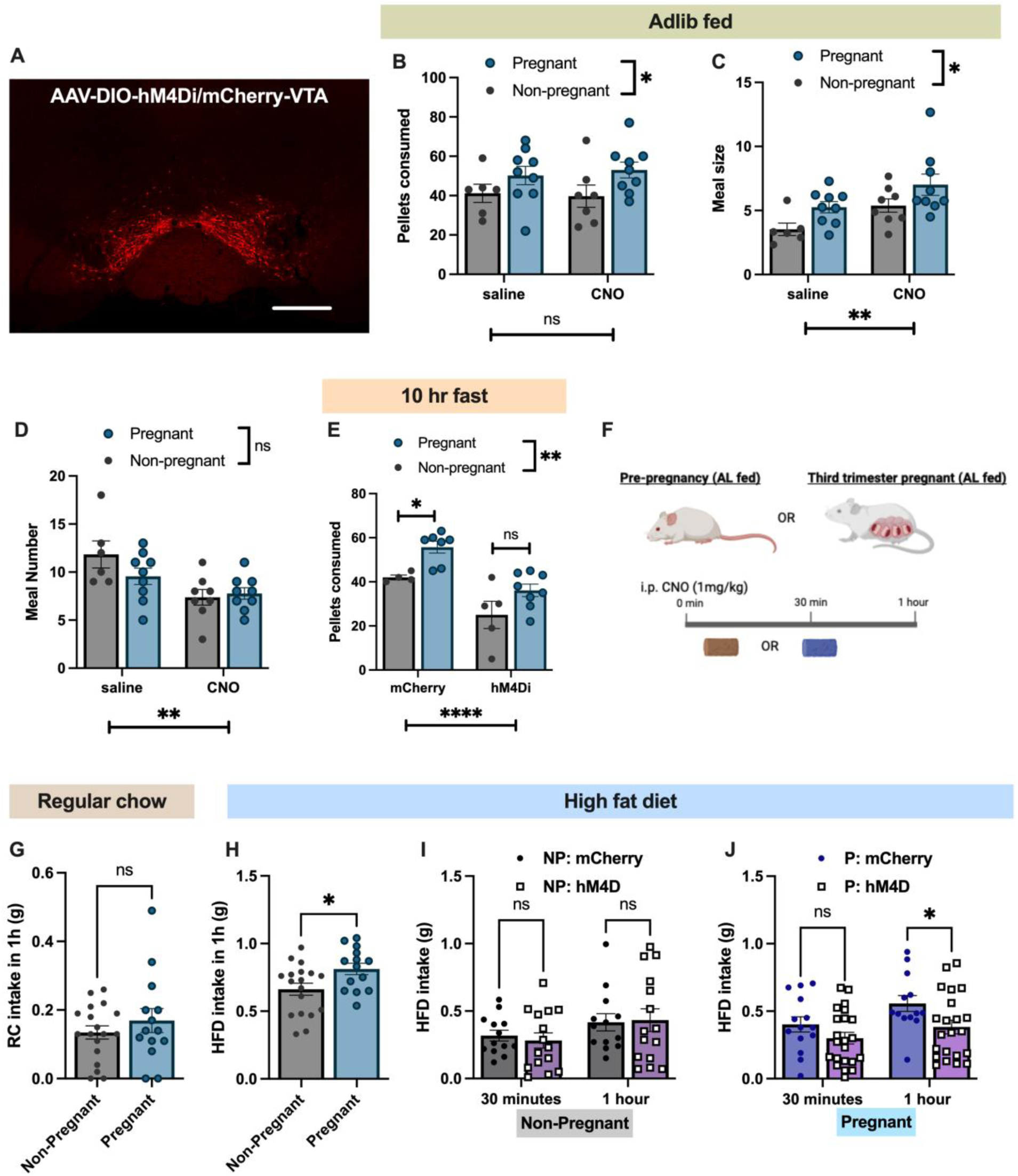
VTA dopamine neurons contribute to increased hedonic feeding and increased sensitivity to negative energy balance in pregnant mice. (A) Representative image of hM4Di-mCherry expression in the VTA. (B) Pellet consumption in non-pregnant and pregnant mice expressing hM4Di-mCherry in VTA dopamine neurons following i.p. injections of saline or CNO. (C and D) Meal size (C) and number of meals (D) following saline or CNO injections in non-pregnant and pregnant mice expressing hM4Di-mCherry in VTA dopamine neurons. (E) Pellets consumed in non-pregnant and pregnant mice expressing either hM4Di-mCherry or control mCherry virus in VTA dopamine neurons following a ten hour fast. All mice were administered CNO ten minutes prior to testing for the experiments shown in E. (F) Schmatic showing experimental design for experiments shown in panels G-J. (G) Regular chow food intake in one hour in non-pregnant and pregnant mice. Although pregnant mice consume more daily calories than non-pregnant mice, no significant difference in 1-hour standard chow food intake is detectable between non-pregnant and pregnant mice during the light period (i.e. during the timepoints in which acute HFD intake was measured in H-J). (H) 1 hour intake of high fat diet in non-pregnant and pregnant mice. (I) High fat diet intake in non-pregnant mice expressing either control mCherry virus or hM4Di-mCherry in VTA dopamine neurons following injections of CNO. (J) High fat diet intake in pregnant mice expressing either control mCherry virus or hM4Di-mCherry in VTA dopamine neurons following injections of CNO. Data points represent individual mice. Scale bar in A (300um). Data in B, C, D, E, I, and J analyzed by 2-way ANOVA. Data in panels G and H analyzed with unpaired Student’s t-test. ns (not significant), *p<0.05, **p<0.01, ****p<0.001.

### VTA dopamine neurons regulate food seeking in pregnant and non-pregnant energy deprived mice

VTA dopamine neurons exhibit increased activity following food consumption in food deprived pregnant mice (**Fig. 2**). To test if VTA dopamine activity contributes to increased food seeking in fasted mice during pregnancy, we measured food intake in non-pregnant and pregnant mice following a ten hour fast. Mice expressing either control mCherry virus or hM4Di in VTA dopamine neurons were administered CNO (1mg/kg) immediately prior to food presentation after a ten hour fast. After an acute fast, pregnant mice consumed more pellets than non-pregnant mice following CNO administration in animals expressing control mCherry virus in VTA dopamine neurons (**Fig. 4E**). However, this effect was attenuated in mice expressing hM4Di in VTA dopamine neurons, such that food intake levels were not significantly different between pregnant and non-pregnant mice during inhibition of VTA dopamine neurons (**Fig. 4E**). In contrast to *ad libitum* fed conditions (**Fig. 4B**), inhibition of VTA dopamine neurons significantly reduced food intake in both non-pregnant and pregnant energy deprived mice, suggesting that VTA dopamine activity is specifically involved in promoting food seeking for regular chow during conditions of negative energy balance.

### VTA dopamine neurons modulate palatable food intake in pregnant mice

Dopamine levels in the nucleus accumbens are elevated during palatable food consumption (**Fig. 3D**), and pregnant mice exhibit increased VTA dopamine activity during palatable food consumption compared to non-pregnant mice (**Fig. 2D**). We thus hypothesized that increased VTA dopamine activity may promote excessive palatable food intake in pregnant mice. To measure acute changes in palatable food intake in non-pregnant and pregnant mice we provided non-pregnant and pregnant mice with standard chow or palatable high fat diet and measured food intake 1-hour later (**Fig. 4F-H**). Although we were unable to detect increased intake of regular chow in pregnant mice during this acute time-period (**Fig. 4G**), consistent with prior acute PB chip feeding assays (**Fig. 1A**), pregnant mice consumed significantly more HFD than non-pregnant mice in acute HFD feeding assays (**Fig. 4H**). To test if VTA dopamine activity is involved in promoting palatable food intake in pregnant mice, we measured acute HFD intake in non-pregnant and pregnant mice expressing either control mCherry virus or hM4Di in VTA dopamine neurons (**Fig. 4I-J**). Inhibition of VTA dopamine neurons did not significantly alter HFD intake in non-pregnant mice (**Fig. 4I**). In contrast, chemogenetic inhibition of VTA dopamine neurons significantly reduced high fat diet intake in pregnant mice (**Fig. 4J).** Thus, VTA dopamine neurons specifically regulate the consumption of palatable high fat diet in pregnant mice.

## Discussion

The pregnancy period requires significant increases in energy intake to accommodate the metabolic demands associated with fetal growth and development^1,2^. Although pregnancy is known to increase feeding behavior, the specific neural circuitry and molecular mechanisms mediating increased feeding during pregnancy are largely unknown. Further, the behavioral mechanisms leading to increased caloric intake during pregnancy (i.e. homeostatic vs hedonic feeding, food seeking behavior, etc) are incompletely understood. Results presented here are consistent with previous findings showing approximately 25% more food intake during the third trimester of pregnancy compared to age-matched non-pregnant mice^1,21,25^. Although pregnant mice overconsume palatable food when provided acute access to high fat diets (**Fig. 1A and Fig. 4H**), pregnant mice adapt to daily HFD access in a similar manner as non-pregnant animals and continue to consume approximately 25% more calories than non-pregnant mice when provided *ad libitum* access to regular chow or HFD (**Fig. 1B and C**).

Despite consuming more food than non-pregnant mice during free access feeding paradigms (**Fig. 1B**), pregnant mice did not demonstrate increased food consumption during both FR3 and FR5 operant food seeking assays. Thus, requiring additional work to obtain food reduced caloric intake in pregnant mice to a level which was indistinguishable from non-pregnant animals. These findings are reminiscent of recent work demonstrating a reduced propensity to diet induced obesity in mice when low levels of work are required to obtain palatable high calorie food (i.e. FR1 or FR3 operant food seeking tasks)^26^. Thus, requiring additional work to obtain food reduces hyperphagia in multiple physiological states associated with positive energy balance (i.e. diet induced obesity and pregnancy). Interestingly, pregnant mice consistently consume larger meals, while eating less frequently during FR3 and FR5 operant assays (**Fig. 1**). This behavioral approach may provide a strategy to maximize energy intake while reducing the amount of time spent foraging for food. Such a behavioral strategy may be particularly useful for pregnant animals, as pregnancy drastically reduces locomotor activity in rodents^25,27^, and thus likely requires adaptations in meal structure to meet the elevated energy demands associated with this period.

Although pregnant mice exhibit increased meal sizes and reduced meal frequency (**Fig. 1**), these changes are unlikely to be mediated by ventral tegmental area dopamine neurons since pregnant mice continued to consume larger meals in the absence of VTA dopamine activity (**Fig. 4C**). However, inhibition of VTA dopamine neurons did overall increase meal size, with similar effects observed in both non-pregnant and pregnant mice (**Fig. 4C**). These findings are consistent with recent work demonstrating a role for VTA dopamine neurons in regulating the size of ongoing meals^28^. Further work is required to determine the neural circuitry and molecular mechanisms mediating altered meal structure in pregnant mice.

Since dopaminergic circuits are important for promoting food seeking and palatable food intake^9^, we hypothesized that mesolimbic dopamine signaling may be altered during pregnancy to promote increased feeding behavior. In vivo imaging data presented here is consistent with this hypothesis as VTA dopamine neurons are hypersensitive to the consumption of palatable food in pregnant mice (**Fig. 2C and D**). Consistently, downstream dopamine levels in the nucleus accumbens are significantly greater in pregnant mice than non-pregnant animals during high fat diet consumption (**Fig. 3C and D**). Therefore, the evoked activity of the mesolimbic dopamine circuitry is increased during palatable food consumption in pregnant mice. Such a response is consistent with increased acute intake of high fat diets in pregnant animals (**Fig. 1A and Fig. 4H**). Ultimately, this elevated dopamine signal is likely mechanistically important for promoting increased acute palatable food intake during pregnancy since chemogenetic inhibition of VTA dopamine neurons reduces high fat diet intake in pregnant mice, but not in control non-pregnant animals (**Fig. 4I, J**). These findings are consistent with a recent report demonstrating that palatable food intake cravings during pregnancy are mediated by increased engagement of D2 dopamine receptors in the nucleus accumbens^29^. Further work is ultimately required to determine the neurophysiological mechanisms mediating increased mesolimbic dopamine activity during pregnancy. Since pregnancy robustly increases the levels of key neuroendocrine hormones (i.e. estrogen and progesterone) which regulate dopaminergic activity^2,30^, future studies are also warranted to map the effect of pregnancy-related hormones on the activity of the mesolimbic dopamine system.

In addition to palatable food intake, mesolimbic dopamine circuitry promotes increased food seeking in energy deprived animals^9,13^. VTA dopamine neurons express receptors for the hunger hormone ghrelin^14,31,32^, which directly activates these cells, and the satiety hormone leptin^12,31,33,34^, which inhibits these cells. Further, acute signals of energy sufficiency such as amylin and glucagon-like-peptide 1 regulate the activity of mesolimbic dopamine circuitry^35,36^. Thus, VTA dopamine neurons are well suited to directly respond to signals of energy availability, linking energy state with food seeking behavior and hedonic feeding^11,16,31,37,38^. Consistent with this hypothesis, both food deprivation and artificial stimulation of hunger via activation of hypothalamic agouti-related peptide (AgRP) neurons increase dopamine levels in the nucleus accumbens during feeding^16,39,40^. Data presented here suggest that pregnancy increases the sensitivity of the mesolimbic dopamine system to energy deprivation. For example, VTA dopamine neuron activity is increased in pregnant mice compared to non-pregnant animals in response to food consumption following a 10-hour fast (**Fig. 2F-H**). Similarly, nucleus accumbens dopamine levels are also elevated in pregnant mice (compared to non-pregnant animals) following an acute fast (**Fig. 3F and G**). Although chemogenetic inhibition of VTA dopamine neurons does not alter regular chow food intake during *ad libitum* fed conditions (**Fig. 4B**), inhibition of these neurons is highly effective at reducing food intake following an acute 10 hour fast (**Fig. 4E**). This effect occurs in both non-pregnant and pregnant mice, but the magnitude of the effect is larger in pregnant animals (**Fig. 4E**). In conjunction with the *in vivo* imaging data presented here, these findings suggest that pregnancy shifts the sensitivity of VTA dopamine neurons to signals of negative energy balance. Further work is required to determine the neurophysiological mechanisms mediating altered VTA dopamine responsivity to hunger signals in pregnant mice, which may involve changes in pregnancy related hormones and/or sensitivity to neuroendocrine signals of energy availability.

In conclusion, the *in vivo* activity of the mesolimbic dopamine system is enhanced in pregnant mice to increase hedonic food intake and the sensitivity of mice to negative energy balance. Given that overfeeding during pregnancy increases the risk of both the mother and her children developing metabolic disorders later in life, our findings suggest that therapies targeting the mesolimbic dopamine system may provide novel pathways to prevent overconsumption during pregnancy.

## Methods

### Animals

All experiments were approved by the University of Illinois Institutional Animal Care and Use Committee (IACUC). Experiments were performed on female mice (8-16 weeks old). Experiments were performed on C57BL6J (Jax#000664) or DAT-Cre mice (Jax# 020080). DAT-Cre were bred in house by breeding Cre heterozygous mice with C57BL6J mice. Litters were genotyped in house with standard PCR primers for the Cre gene to confirm the transgenic allele: Cre common (5’ GCT TCT TCA ATG CCT TTT GC 3’) and Cre mutant (5’ AGG AAC TGC TTC CTT CAC GA 3’). Prior to experiments mice were group housed in 2-5 mice per cage, in a temperature (20 C) and humidity-controlled environment, with a 12 h light/dark cycle. *Ad libitum* access to food and water was always provided unless specifically mentioned in the text (i.e. during fasting experiments). To generate pregnant and control mice for experiments, mice were bred to a reproductively experienced male mouse for 5 days. Mice were checked daily for a vaginal plug indicating successful mating, which was marked as pregnancy day 1. Control virgin mice were instead paired with a female mouse for 5 days. Following mating, all mice were single caged and body weight and food intake was measured to confirm hyperphagia and weight gain in pregnant mice. In animals subjected to surgical procedures, animal breeding occurred between 2-4 weeks following viral injections. All experiments were performed during the third trimester (day 14-20) of pregnancy or the equivalent time-period in non-pregnant control mice. Dopamine fiber photometry and feeding behavior assays were performed on C57/BL6J mice purchased from Jackson Labs that were approximately matched for age. For experiments involving transgenic mice (i.e. DAT-Cre mice), littermates were used as control animals.

### Viral Vectors

Adeno-associated viral vectors (AAV) that were used in this study included Cre-dependent GCAMP6s (AAV5-Syn-Flex-GCAMP6s-WPRE-SV40; #100845), Cre-dependent hM4Di (AAV5-hsyn-DIO-hM4Di-mCherry; #44362), Cre-dependent mCherry control virus (AAV5-hsyn-DIO-mCherry; #50459) or Cre-dependent GFP control virus (AAV5-hsyn-DIO-EGFP; #50457), and GRAB dopamine sensor virus (AAV9-hsyn-GRAB-DA2m; #140553). All viruses were purchased from addgene and were injected into the brain at stock concentrations (> 1×10^12 vg/mL).

### Stereotaxic viral injections and fiber placements

Stereotaxic surgeries were performed as described in our prior studies^41^. Mice were anesthetized with isoflurane and placed in a stereotaxic apparatus (Kopf) with a constant flow of oxygen and isoflurane during surgeries. Mice were administered preoperative carprofen (5mg/kg, s.c.) for pain prior to injections and for two days post surgeries. To inject virus and/or implant fiber optic inplants, a small incision was made on the skull in the area between bregma and lambda. AAV vectors were injected into the nucleus accumbens or ventral tegmental area using a pulled glass micropipette, which was attached to a micromanipulator (Ronal Tool). Viral injection coordinates for targeting the ventral tegmental area with GCAMP6s (300nl) or hM4Di/mCherry (200nl) virus were as follows (from bregma): A/P: -2.8 and -3.1mm, M/L: +/-0.35mm, D/V: -4.2mm (from the surface of the brain). GRAB dopamine sensors were injected into the nucleus accumbens at the following coordinates: A/P:1.2mm, M/L=-1.0mm: D/V=-3.5mm, -3.8mm and -4.1mm. Injections of hM4Di/mCherry were bilateral while injections of dopamine GRAB sensors or GCAMP6s was unilateral. For each injection, virus was injected over 5-10 minutes and left for an additional 5 minutes before removing the needle.

For fiber photometry experiments, during the same surgery, a fiber optic cannula (200um, RWD Biosciences) was implanted directly about the ventral tegmental area or nucleus accumbens viral injection sites. After fiber insertion, the fiber was secured to the skull using dental cement (C&B Metabond). Following surgeries, mice were single caged and returned to housing facilities for at least three weeks before starting experiments.

### Fiber photometry experiments

Fiber photometry equipment and analysis was performed as described in our prior study^41^ Mice were connected to a Plexon Multi-Wavelength Fiber Photometry System (Plexon, 8-61-A-07-A) via a fiber optic patch cord (Plexon, 08-60-A-04-C). Patch cords were attached to the fiber optic implant on the mouse’s head via mating sleeves (Plexon). Blue (465nm) and UV (410nm) light sources were provided by internal LED drivers which are built into the Plexon fiber photometry system. Fluorescent signals are recorded by the Plexon photometry system which cycles on and off at 30hz sampling window between the 410nm (isosbestic control signal) and 465nm (GCAMP6 signal) signals. Analysis of fiber photometry analysis was performed using a custom R code, as described in our prior study^41^

After allowing 2-4 weeks for viral expression and recovery from surgery, all mice were first tested for viral expression by recording the responsivity (for both VTA GCAMP and NAc GRAB sensors) of sensors to acute consumption of a high fat diet pellet. Animals only had access to this diet for this initial 10-minute test. Only animals with noticeable increases in signal in response to HFD were included in subsequent experiments. All mice were subsequently tested in two behavioral assays, which were performed both prior to pregnancy (baseline period) and during the third trimester of pregnancy (or the equivalent time-period in non-pregnant control mice). First, we tested the response of nucleus accumbens dopamine to presentation of a standard chow food pellet in the fed state. Following ten minutes of baseline recording, a food pellet was presented to all mice for an additional five minutes. Following this experiment, on the same day, mice were disconnected from the fiber setup and returned to their cages for an additional ten hours. Food was removed from all cages during this ten-hour period to test the response to chow presentation in energy deprived mice (food removed from 7am-5pm). The identical experiment was subsequently performed on the same mice following a ten hour fast. 2-4 days later, the same mice were tested for their response to consumption of palatable high fat diet in the fed state. For these experiments, all mice were habituated to the high fat diet by providing access to HFD in the home cage for ten minutes for two days prior to the photometry experiment. On the experimental day (experiment performed during the light period: between 10am-4pm), mice were again attached to the fiber photometry system and baseline signal was recorded for ten minutes. Following baseline recordings, mice were provided with high fat diet for an additional ten minutes. For both fast-refeeding and HFD photometry assays a plexon event input generator was used to manually score the start of eating, and these events were directly aligned with fiber photometry calcium traces for analysis. Following this initial testing, all mice were approximately divided into “pregnant” and “non-pregnant” groups. The pregnant group was bred to reproductively experienced male mice following baseline experiments, while the non-pregnant group was paired with a female mouse. The same two experiments (10 hr fast and high fat diet presentation) were again repeated on the same mice during either the third trimester of pregnancy or the equivalent time-period in the non-pregnant control group. Changes in calcium response to events (high fat diet consumption or regular chow consumption) were calculated in the 60- or 180-seconds following events (i.e. start of food consumption) compared to the 60 seconds prior to each event (i.e. prior to food consumption). The change in calcium signal was calculated for each mouse during the two time points in to determine the effect of pregnancy on nucleus accumbens dopamine responses (within subjects’ comparison). We also directly compared the response between pregnant and non-pregnant mice on the same test day (i.e. between subjects’ comparison). Although we observed stable recordings of dopamine sensor recordings at each time-point in the control non-pregnant mice (i.e. comparing time point 1 to time point 2), mice expressing GCAMP6 in VTA dopamine neurons exhibited signal decay between the two time-points. Therefore, for GCAMP6 experiments we only compared the change in calcium signal in non-pregnant vs pregnant mice during the same testing day (i.e. between subjects’ comparisons).

### Feeding behavioral assays and experiments with FED3 devices

All feeding experiments were performed with feeding experimental devices, except for high fat diet measurements (and initial characterization of regular chow and high fat diet intake in non-pregnant and pregnant mice (i.e. Figure 1A and B)), which were manually measured in the mouse home cage. For manual food intake measurements, a pre-measured amount of food was added to the mouses home cage and the change in the weight of the food was measured the following day. Cages were changed daily during testing to prevent spillage.

FED3 feeding assays were performed as described in our prior study^41^. Briefly, FED3 devices were attached to the side of the mouse’s cage, and mice obtained all their food from FED3 devices. Fixed Ratio 1 (FR1) schedule of reinforcement, in which one nose poke on the left poke resulted in dispensing of one 20mg food pellet, were utilized for all feeding assays, except when FR3 and FR5 assays are specially mentioned in the text and figure legends (i.e. Figure 1). We began collecting experimental data after mice had reached at least 70% correct nose pokes (i.e. 70% of nose pokes occur on the correct left poke vs the incorrect right poke), typically 1 -2 days following the start of testing. FR1 was chosen over free access feeding as we observed significantly less food hoarding in FR1 schedule vs free feeding mode during prolonged feeding measurements. For all schedules of reinforcement, feeding patterns were characterized by meal size and meal frequency. We defined a meal as the number of pellets taken with an inter-pellet interval between each pellet less than 5 minutes, while meal frequency was the number of meals in 24 hours. Data was collected every day at ZT6.

For regular chow chemogenetic feeding experiments with the FED3 devices (Fig. 4), mice were administered saline (i.p., 200ul) or CNO (1mg/kg, i.p.) in a randomized fashion during the light period. Changes in pellets consumed, meal size, and meal number were calculated for each mouse following saline or CNO injections and compared for statistical analysis (repeated measures comparison). For HFD chemogenetic assays, all mice received HFD for ten minutes in the two days prior to testing to habituate to HFD presentation. On the testing day, all mice (both mice expressing control mCherry or hM4Di in VTA dopamine neurons) were administered CNO (1mg/kg) fifteen minutes prior to providing high fat diet to the mice. Consumption of high fat diet was measured 30 minutes and 1 hour following i.p. injections and compared between the mCherry and hM4Di expressing mice. For fasting experiments, after mice were trained on FR1 assays with FED3 devices, all mice were fasted for ten hours during the light period (7am-5pm). Following fasting, all mice were administered CNO (1mg/kg) ten minutes prior to providing FED3 devices to the mice.

Experiments involved DREADD targeting of VTA dopamine neurons were peformed on two cohorts of mice. High fat diet feeding experiments were performed on both separate cohorts of mice, and combined data from both cohorts are plotted in figure 4. Fasting experiments were only performed on one cohort of animals. Representative viral locations from all mice for both cohorts are shown in extended figure 2.

### Post-hoc validation of viral and fiber optic placement

Following the completion of behavioral experiments on virally targeted and fiber implanted mice, all mice underwent trans-cardiac perfused to check for the location of virus and fiber optic placement. Perfusions and post-hoc fixation/cryopreservation were performed as described in our prior study^41^. Brain sections covering the VTA or NAc were obtained in 40um sections using a cryostat (Leica CM3500) and mounted onto glass slides to check for viral location and fiber optic placement. Only mice with correct targeting of virus into the VTA/NAc and fiber optic placement in VTA/NAc were included in experiments.

### Statistical Analysis

Specific statistical tests are outlined in the figure legends. Data that was normally distributed was analyzed with parametric statistical tests, while data that was not normally distributed was analyzed with non-parametric tests. Data was analyzed using Graphpad Prism.

### Funding Sources

This work was funded by the University of Illinois and the National Institute of Health (R01HD113522 to PS)

---

**Extended Data Fig. 1:**
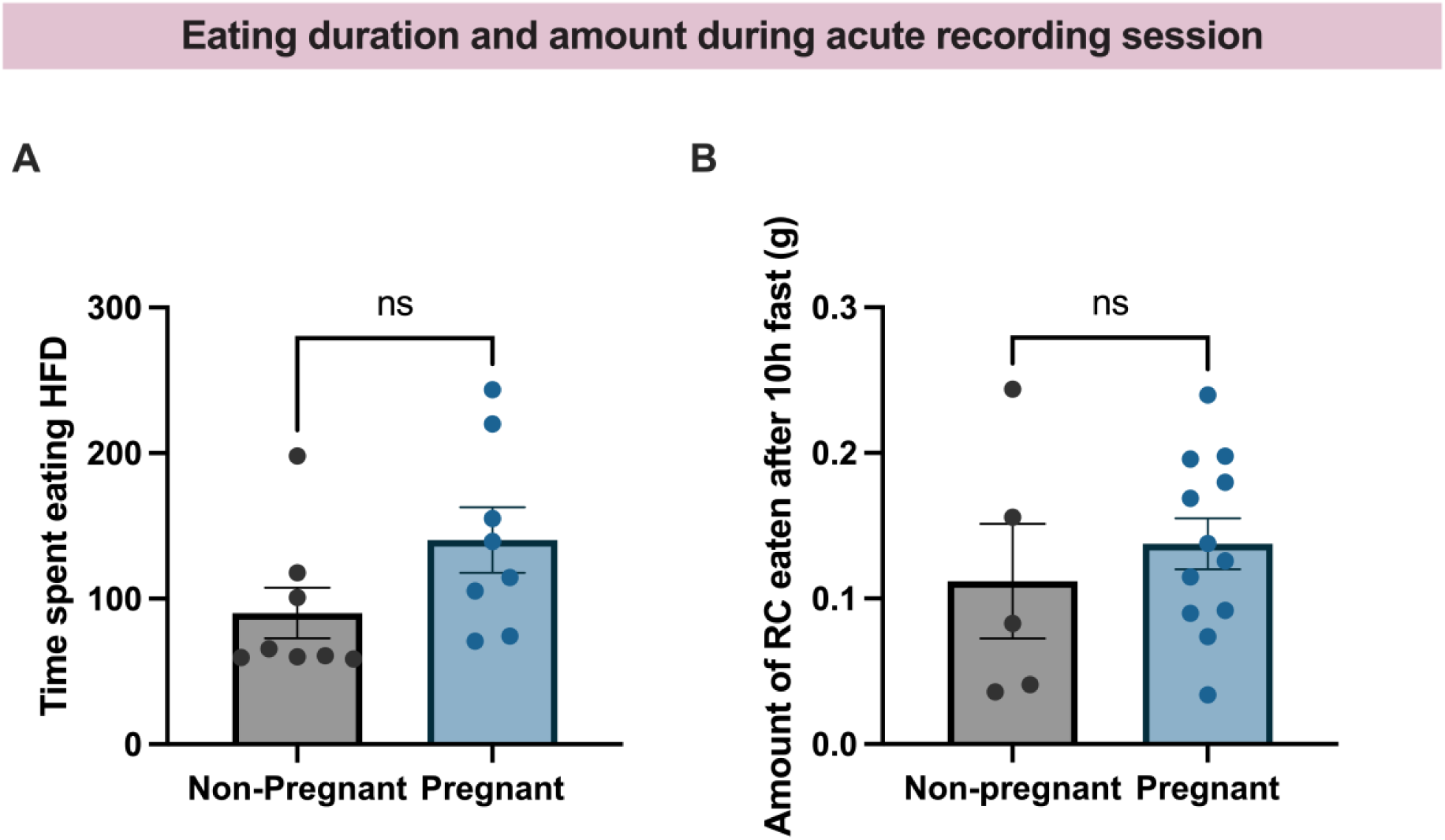
Time eating does not differ between pregnant and non-pregnant mice during fiber photometry recordings of VTA GCAMP signal. (A) Time spent eating the high-fat diet by the non-pregnant and pregnant mice in the 10-minute fiber photometry recording session. (B) Amount of regular chow eaten by non-pregnant and pregnant mice during the first 10 mins of refeeding. Data points represented individual mice. Data represented as mean ± SEM. Data analyzed using unpaired student’s t test.

**Extended Data Fig. 2:**
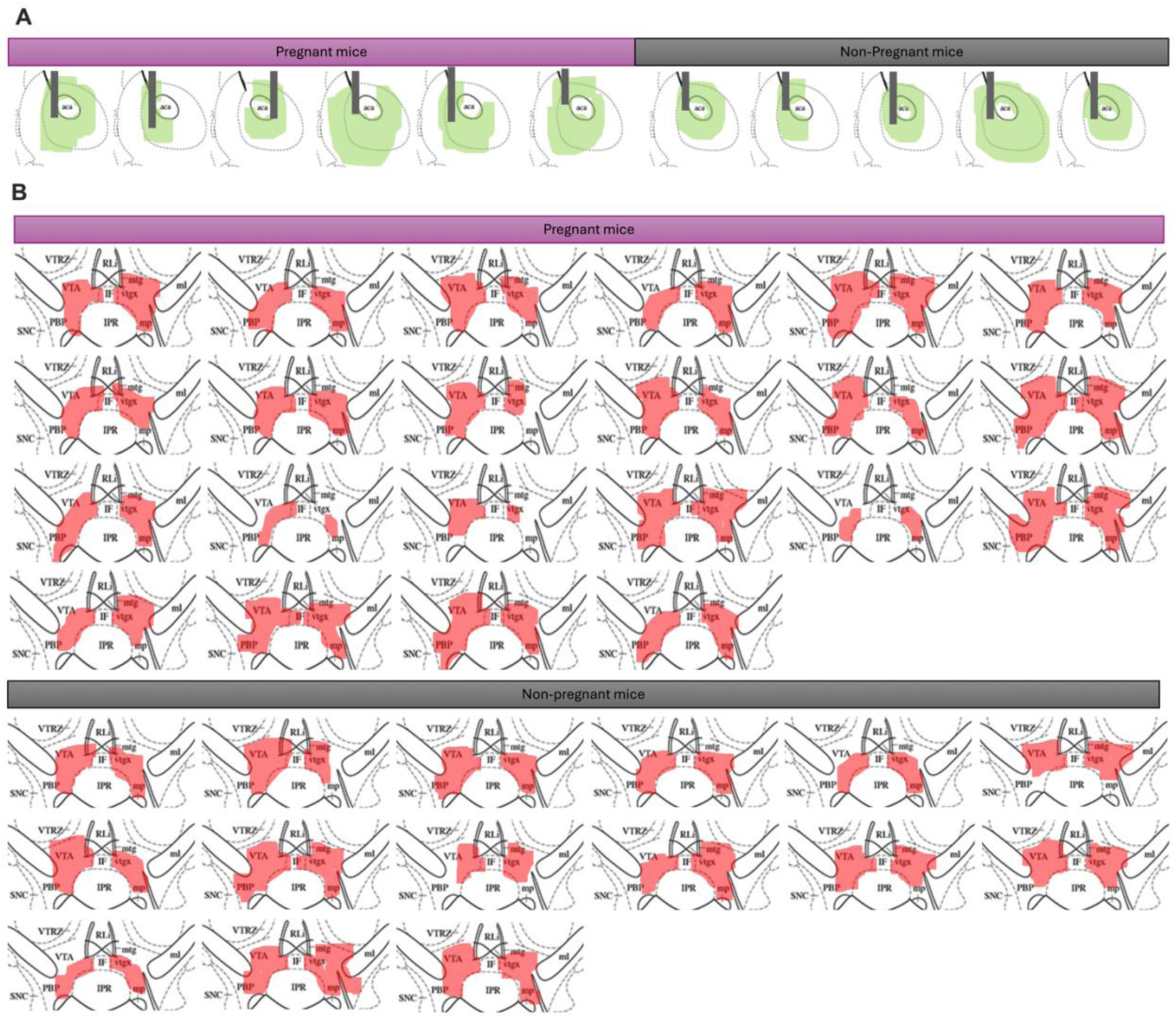
Representative viral locations and fiber optic placements for nucleus accumbens GRAB dopamine imaging experiments. (A) Schematics showing the location of the viral GRAB sensor and fiber location in the pregnant and non-pregnant mice for GRAB fiber photometry experiments. Spread of the virus is shown in green, while location of fiber optic is shown as grey line. (B) Schematics showing the spread of viral expression of the inhibitory DREADD in the pregnant and non-pregnant mice for chemogenetic studies. The spread of the virus in the VTA is shown in red.

**Extended Data Fig. 3:**
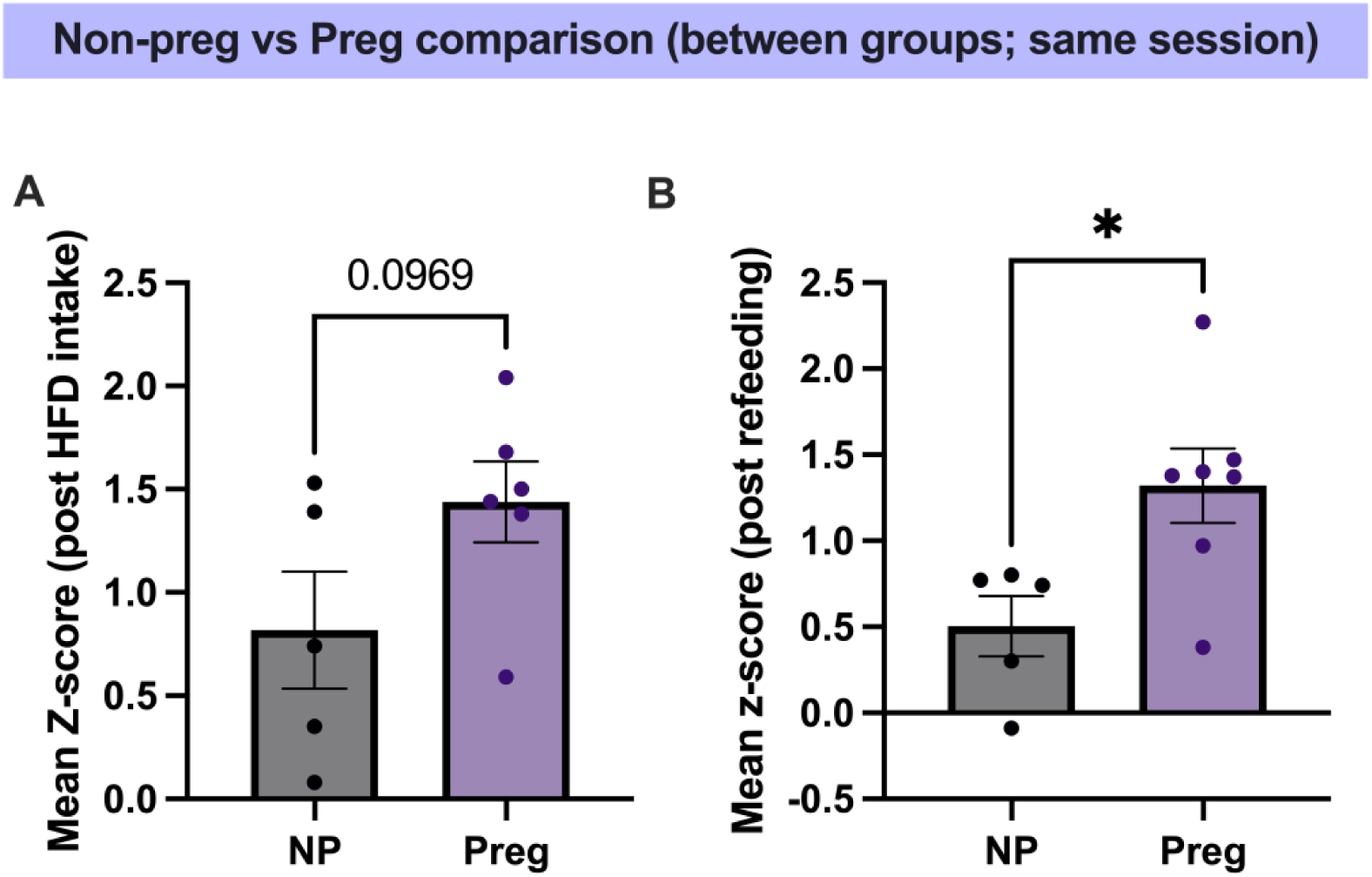
Nucleus accumbens dopamine signal is enhanced in pregnant mice during homeostatic and hedonic feeding tasks. (A) Comparison of average z-score during consumption of HFD in pregnant and non-pregnant mice in the same session. (B) Comparison of the average z-score while refeeding in pregnant and non-pregnant mice in the same session. Data points represented individual mice. Data represented as mean ± SEM. Data analyzed using unpaired student’s t test.

**Extended Data Fig. 4:**
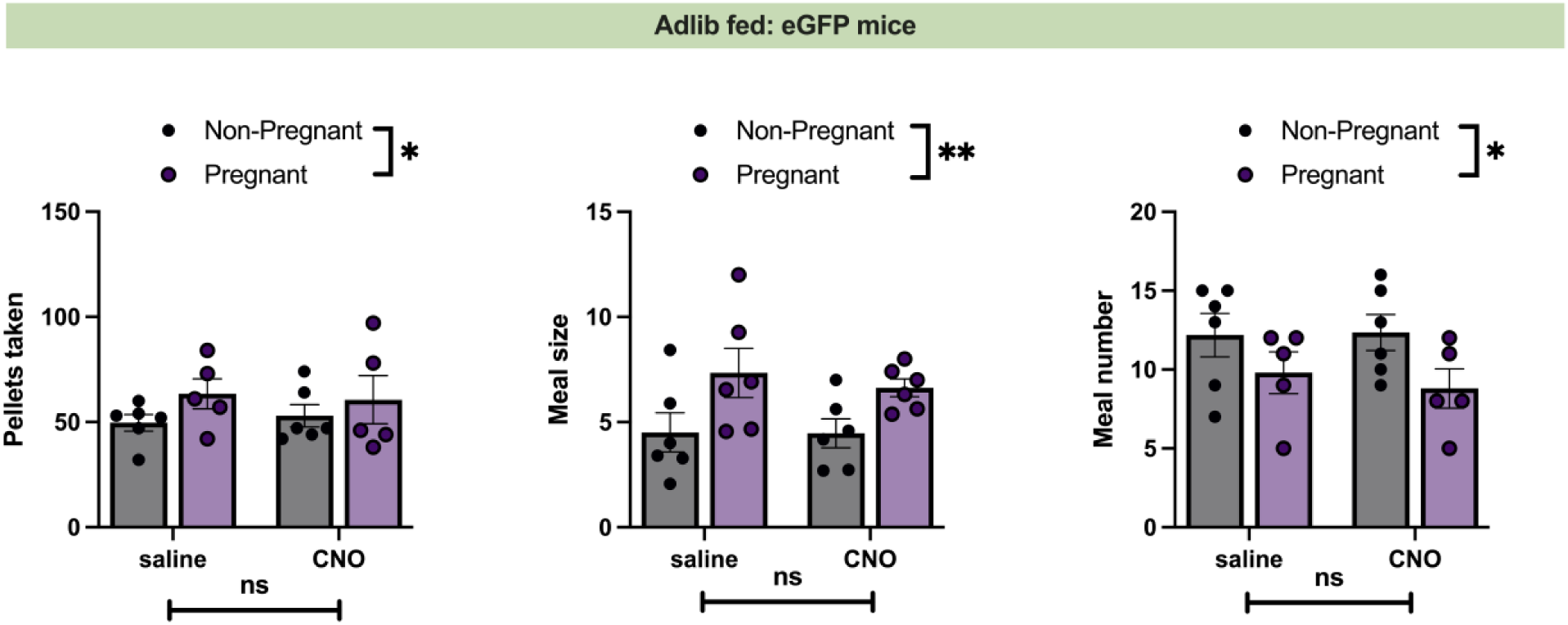
CNO administration does not alter feeding behavior in control mice expressing eGFP in VTA dopamine neurons. (Left) Number of pellets taken in 4 hours following administration of either saline or CNO in control eGFP injected non-pregnant and pregnant mice. (Middle) Average meal size in the 4 hours following administration of either saline or CNO in the eGFP injected non-pregnant and pregnant mice. (Right) Number of meals in the 4 hours following administration of either saline or CNO in the eGFP injected non-pregnant and pregnant mice. Data represented as mean ± SEM. Data points represent individual mice. Data analyzed by 2-way ANOVA. *p<0.05, **p<0.01, ns (not significant).

